# A systematic comparison of error correction enzymes by next-generation sequencing

**DOI:** 10.1101/100685

**Authors:** Nathan B. Lubock, Di Zhang, George M. Church, Sriram Kosuri

## Abstract

Gene synthesis, the process of assembling gene-length fragments from shorter groups of oligonucleotides (oligos), is becoming an increasingly important tool in molecular and synthetic biology. The length, quality, and cost of gene synthesis is limited by errors produced during oligo synthesis and subsequent assembly. Enzymatic error correction methods are cost-effective means to ameliorate errors in gene synthesis. Previous analyses of these methods relied on cloning and Sanger sequencing to evaluate their efficiencies, limiting quantitative assessment and throughput. Here we develop a method to quantify errors in synthetic DNA by next-generation sequencing. We analyzed errors in a model gene assembly and systematically compared six different error correction enzymes across 11 conditions. We find that ErrASE and T7 Endonuclease I are the most effective at decreasing average error rates (up to 5.8-fold relative to the input), whereas MutS is the best for increasing the number of perfect assemblies (up to 25.2-fold). We are able to quantify differential specificities such as ErrASE preferentially corrects C/G → G/C transversions whereas T7 Endonuclease I preferentially corrects A/T → T/A transversions. More generally, this experimental and computational pipeline is a fast, scalable, and extensible way to analyze errors in gene assemblies, to profile error correction methods, and to benchmark DNA synthesis methods.

## Introduction

Synthetic DNA is a central tool for biological research [1]. Notably, the initial development of nucleic acid synthesis led directly to the cracking of the genetic code [2]. Today, progress in biology is often limited by the difficulty in producing long, high-quality synthetic DNA [3, 4]. This bottleneck is particularly apparent in the assembly of gene-sized fragments of DNA known as gene synthesis [5]. Currently, gene synthesis relies on the assembly of many oligonucleotides (oligos) of ∼40-150 nucleotide (nt) into a single larger piece of DNA of >1,000 base-pairs (bp) [5]. A variety of methods to assemble oligos into gene-sized fragments exist, but ligation- and polymerase-based assembly methods are the most common [6, 7, 8, 9]. Regardless of the method, the quality of the final product is largely dependent on the quality of the oligos used in the assembly.

Oligos are primarily synthesized using phosphoramidite chemistry first developed by Beaucage and Caruthers in the 1980s [10]. Although these oligos are of high enough quality for common applications such as PCR, their error rates make practical gene synthesis challenging. Several groups have managed to synthesize genes from such oligos, but only find about 5-60% perfect products depending on the size and complexity of the template [11, 12, 13, 14]. This problem is further exacerbated when using lower-cost, but often lower quality oligos from array-based synthesis approaches [15, 16, 17, 18, 19, 20].

Consequently, researchers have developed a number of methods to ameliorate oligo error rate post-synthesis. Size selection methods such as HPLC or PAGE can filter truncated sequences, but are labor-intensive and ineffective against small errors such as single-base deletions, insertions, or substitutions [21, 22]. Hybridization-selection techniques can filter large pools of oligos, but are cost-prohibitive as the number of oligos needed effectively doubles [16, 23]. Sequencing-based retrieval methods can physically pick perfect sequences or separate them by barcoded PCR, but are time-intensive and can require specialized equipment [24, 25, 26]. Enzymatic error correction is a more commonly-used technique that is relatively inexpensive and effective against most errors. This method employs a variety of different enzymes traditionally used for mutation detection to filter out by binding to or cutting at errors [27, 28, 29, 30].

Two particular classes of proteins are most prevalent in error correction: mismatch binding proteins and mismatch cleaving proteins. Generally, these enzymes recognize distortions in the DNA helix that are caused by mishybridized bases on either strand. In gene synthesis, a pool of perfect and imperfect sequences will be melted and re-annealed pairing perfect and imperfect strands to one another. This produces mishybridized bases that can be recognized by these enzymes. Mismatch binding proteins are used to enrich perfect sequences, while mismatch cleaving proteins are used (often in conjunction with exonuclease trimming) to remove imperfect sequences. The most commonly used mismatch binding protein, MutS, recognizes and binds to all single-base mismatches and a variety of small single stranded loops caused by insertions or deletions (indels) with varying affinity [31, 32, 33, 34, 35]. There are a number of different ways to bind and separate error-containing DNA with MutS including: gel-shift assays, MutS-functionalized columns, and MutS-functionalized magnetic beads [11, 20, 36]. Mismatch cleaving enzymes operate by cutting at or near an error and a variety of different mismatch cleaving enzymes are in use [37]. Broadly, these enzymes can correct errors in two different ways. Similar to mismatch binding methods, perfect sequences can be recovered by filtering them from those cut by mismatch cleaving enzymes. Alternatively, the exonuclease activity is used to trim the error-containing region left over by the mismatch cleaving enzymes. The full length sequences are then recovered by performing a PCR assembly with the trimmed sequences.

Previous assessments of different enzymatic error correction methods have relied on Sanger sequencing of finished gene synthesis products to determine their efficiencies [11, 12, 14, 19, 20]. These studies find that, broadly, the dominant mode of errors in gene synthesis products are single-base deletions and mismatches. However, the prohibitive cost of Sanger sequencing hundreds of thousands of bases has limited the effective characterization and comparison of existing methods. Alternatively, one can turn to the mutation detection literature to find biochemical characterizations of enzymes commonly used in error correction [30, 34, 38, 39, 40]. Although these reports provide more detailed affinity data, they typically rely on electrophoretic methods and are thus similarly limited in sample size.

In order to overcome these limitations, we developed a custom experimental and computational pipeline that leverages Next-generation Sequencing (NGS) to characterize error rates. Here we report the first in-depth characterization via NGS of both the errors arising from the assembly process, as well as the ability of six of the most commonly used error correction enzymes to eliminate these errors across 11 total conditions. With sample sizes three to four orders of magnitude larger than previous reports, we are able to gain detailed insights into the modality of errors as well as each enzyme’s relative ability to correct them. We believe that our method can act as a generalizable platform to rapidly and cost-effectively test, characterize, and optimize oligo synthesis parameters or new enzymatic error correction methods.

## Materials and Methods

### Pre-processing

To ensure that we only analyzed high quality reads, we first ran our sequencing data through a pre-processing pipeline. First, we used `BBDuk` (part of the `BBMap` suite; version 36.14) to trim any Illumina adapters from our reads [41]. Next, we used `BBDuk` to remove any reads with at least 26 bases that match to the PhiX (NC_001422) or *E. coli* (U00096.3) genomes. We also removed any read pairs that had an “N” base call in either one of the reads during this step. We then took the filtered reads and merged read pairs with perfectly overlapping regions with `BBMerge` (also part of the `BBMap` suite; version 36.14) using the `pfilter=1` option.

### Alignment and Parsing

After read pre-processing and merging, we use a custom Python script to align our reads to the reference oligo sequence, and parse the resulting alignments to get the positions of all errors. Our Python script uses the `uta-align` (version 0.1.6) package from the Python Package Index (PyPI) to perform a Needleman-Wunsch exhaustive global alignment of the input reads to the reference sequence [42]. Our script can also provide functionality for performing any alignment supported by the `uta-align` library (e.g. Smith-Waterman local alignments), and allows for tunable gap penalties or match scores.

Once the alignment and parsing is complete, our script will output the results in a tidy csv file with the name of the read, the position of the error, the type of error, and the actual error itself [43]. The types of errors are as follows: M - Mismatch, D - single-base Deletion, I - single-base Insertion, P - multiPle-base deletion, and S - multiple-base inSertion. The errors are classified as: (Original Base)(Mutated Base) for mismatches; the reference base(s) that were deleted for deletions; and the base(s) that were inserted for insertions. Both single and multiple-base insertions are mapped to the “right” of the base in the reference sequence. For example, if the reference sequence was “GATTACA” and we inserted a C at position 3, the resulting alignment can be visualized as:

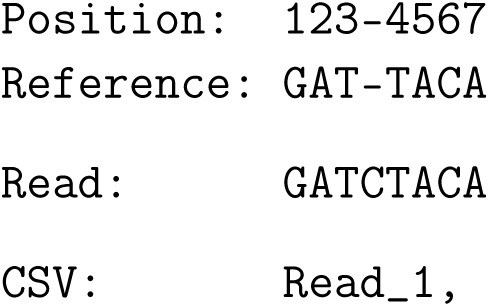

Lastly, if there is a single-base deletion or insertion in a region where there is an identical base adjacent to the mapped position of the error, we distribute the fractional count of the total number of identical bases over each position. For example, if our alignment produced a deletion of A at position 2 in the sequence “TAAAG,” our software will note this as a deletion of A at positions 2, 3, and 4, with fractional counts of 1/3 at each of those positions. This compensates for the fact that there are three equally valid alignments in that region.

### Error Frequency Calculations and Definitions

To be consistent with previous studies, we calculated the relative error frequency per kb (*f*) as

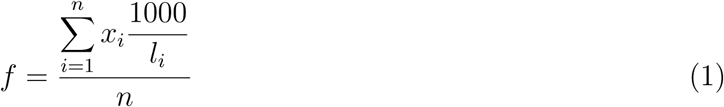

where *x_i_* is the number of errors in read *i*, *l_i_* is the length of that read, and *n* is the total number of reads [12]. This is distinct from error rates, which are defined as the number of errors detected at given base, divided by the total number of sequencing reads in the sample. Error rates can be further separated by the specific error sub-type.

### Reagents

All the oligos were synthesized by Integrated DNA Technologies (IDT). The ErrASE Error Correction Kit was purchased from Novici Biotech and is now available as CorrectASE from Life Technologies. The Surveyor Mutation Detection Kit was from Transgenomic. T4 Endonuclease VII was from Affymetrix. *Thermus aquaticus* MutS DNA mismatch repair protein was from Excellgen. Endonuclease V, T7 Endonuclease I, and T7 DNA Ligase were all from New England Biolabs.

### Error-enriched oligonucleotide synthesis and template assembly

The 85-nucleotide (nt) forward (5’ TACACGACGCTCTTCCGATCTGCTGCCGATTTCCATAAGATGCCTCCACGTCTCCGAAGAACTACATGGTGAATGTGTGAAGGCA 3’) and reverse (5’ AGACGTGTGCTCTTCCGATCTCGATATATACAACACTGCTCGAGGATTGGTTCAAAATGCCTTCACACATTCACCATGTAGTTCT 3’) oligos contains 21nt primer sites and 64nt template regions, 63 of which, except for the last base, were doped with 3% errors at each position. This doping is achieved by hand-mixing 1% of every other base into the 97% of the reference base. For example, according to the reference sequence, if a position is supposed to be an A, then 1% of C, T, and G was mixed into 97% A during the initial oligo synthesis by IDT. With 28nt complementary regions, the two oligos were able to anneal and then assembled into a 142-base pair (bp) doubled-stranded template. This template consists of two 21bp primer regions and a 100bp region (5’ GCTGCCGATTTCCATAAGATGCCTCCACGTCTCCGAAGAACTACATGGTGAATGTGTGAAGGCATTTTGAACCAATCCTCGAGCAGTGTTGTATATATCG 3’) for error correction and for subsequent next-generation sequencing.

Specifically, to pre-assemble the forward and reverse oligos, 10.4*μ*L nuclease-free water (Ambion), 4*μ*L 5X HF Buffer (New England Biolabs), 0.4*μ*L 25mM dNTP (New England Biolabs), and 0.2uL Phusion High Fidelity Polymerase (New England Biolabs) were added into 5*μ*L 1*μ*M mixed aforementioned forward and reverse oligos. Initially heated at 98 °C for 30 seconds, the reaction was then cycled 15 times: at 98 °C for 5 seconds, at 70 °C for 1 second, ramping down with a speed of 0.5 °C/second to 50 °C, at 50 °C for 30 seconds, and at 72 °C for 20 seconds. The final extension step was at 72 °C for 5 minutes. The product after the pre-assembly step was diluted 1:10 in nuclease-free water, 2*μ*L of which, served as template, was added into 35.25*μ*L nuclease-free water, 10*μ*L 5X HF Buffer, 1*μ*L 25mM dNTP, 0.5*μ*L Phusion High Fidelity Polymerase, 1.25*μ*L 10mM mixture of forward (5′ TACACGACGCTCTTCCGATCT 3′) and reverse (5′ AGACGTGTGCTCTTCCGATCT 3′) PCR amplification primers to make the total volume of this PCR 50*μ*L. Initially heated at 98 °C for 30 seconds, the reaction was then cycled 25 times: at 98 °C for 5 seconds, at 62 °C for 10 seconds, at 72 °C for 10 seconds. The final elongation step was at 72 °C for 5 minutes. Pooled PCR products were then cleaned using QIAquick PCR Purification Kit (Qiagen), and the purified products served as the template for subsequent error correction treatments and sequencing.

### Error correction of the synthetic DNA template

#### ErrASE

Per the manufacturer’s instructions, 60*μ*L of ∼50ng/*μ*L template in 1X HF Buffer was re-annealed to form heteroduplex by heating at 98 °C for 1 minute, cooling at 0 °C for 5 minutes, and incubating at 37 °C for 5 minutes. Next, 10*μ*L of this re-annealed heteroduplex was added into each well of the 6-well ErrASE tube and was incubated at room temperature for 1 hour. We then combined 2*μ*L from each well as template into the recovery PCR, whose setup and thermocycling conditions were the same as the assembly PCR in the section above. The PCR product using the treated heteroduplex from the first well of the ErrASE tube (presumably has the highest concentration of ErrASE) presented a band, indicating successful recovery after error correction. This product was thus cleaned-up using QIAquick PCR Purification Kit and served as the template for the second iteration of ErrASE treatment.

#### Surveyor

Per the manufacturer’s instructions, ∼50ng/*μ*L template in 1X HF Buffer was re-annealed to form heteroduplex by the following thermocycling conditions. First, the sample was heated at 95 °C for 10 minutes. Then, the temperature was ramped down at 2 °C/second, and was held at 85 °C for 1 minute. Finally, the temperature was further cooled down to 25 °C at 0.3 °C/second, and was held for 1 minute at every 10 °C interval. Per Saaem *et al.*, 2*μ*L Surveyor Nuclease S and 1*μ*L Enhancer S were added into 8*μ*L re-annealed heteroduplex [19]. The reaction mixture was then incubated at 42 °C for 60 minutes. After the treatment was concluded, 2*μ*L of the mixture served as the template in the recovery PCR, whose setup and thermocycling conditions were the same as the assembly PCR. The product of this recovery PCR, once cleaned-up, entered the next round of Surveyor Nuclease treatment.

#### Endonuclease V

Similar to Fuhrmann *et al.*, 10*μ*L of ~50ng/*μ*L template in 1X HF Buffer was re-annealed using the cycling condition described in the ErrASE section [12]. We then added 5U of Endonuclease V, 2*μ*L of NEBuffer 4, and nuclease-free water to the re-annealed heteroduplex to make the total volume 20*μ*L. The reaction was incubated at 37 °C for 24h, and 2*μ*L of this mixture served as the template for the recovery PCR. The cleaned-up product then entered the next iteration of Endonuclease V treatment.

#### T7 Endonuclease I (Fuhrmann)

As in Fuhrmann *et al.*, 10*μ*L of ∼50ng/*μ*L template in 1X HF Buffer was re-annealed using the cycling condition described in the ErrASE section [12]. We combined 2*μ*L of NEBuffer 2, 25U of T7 Endonuclease I, and nuclease-free water to make the final volume 20*μ*L. The reaction was incubated at 37 °C for 24 hours, and 2*μ*L of the mixture served as the template for the recovery PCR. The cleaned-up product entered the next iteration of T7 Endonuclease I treatment.

#### T7 Endonuclease I with T7 DNA Ligase

We first re-annealed 100ng of template in 1X HF Buffer according to the ErrASE protocol. Then we combined 2.5*μ*L of T4 DNA Ligase reaction buffer, 10U of T7 Endonuclease I, T7 DNA Ligase (at 0, 1000U, or 10000U), and the appropriate amount of nuclease-free water to make the final volume 25*μ*L. The reaction was then incubated at 25 °C for 4 hours, and 2*μ*L of the treated sample served as the template for recovery PCR. We used 100ng of the cleaned-up product for the next iteration of T7 Endonuclease I/T7 DNA Ligase treatment.

#### T4 Endonuclease VII

First, 10*μ*L of ∼50ng/*μ*L template in 1X HF Buffer was re-annealed using the cycling condition described in the ErrASE section. Then, 1*μ*L 1M Tris-HCl (pH 8.0), 4*μ*L 50mM MgCl_2_, 2*μ*L 100mM *β*-mercaptoethanol, 1*μ*L 10mg/ml BSA, and 2*μ*L T4 Endo VII (1000U) was added to the 10uL heteroduplex. The reaction mixture was incubated at 37 °C for 24 hours, and 2*μ*L of which served as the template for the recovery PCR. Then the cleaned-up PCR product entered the next cycle of T4 Endonuclease VII.

#### MutS

Per the manufacturer’s instructions, 250ng/*μ*L in 10mM Tris-HCl (pH=7.8) and 50mM MgCl_2_ was heated to 95 °C for 5 minutes followed by cooling at 0.1 °C/second to 25 °C. To the re-annealed template, 207.39*μ*L 1X binding buffer (20mM Tris-HCl (pH=7.8), 10mM NaCl, 5mM MgCl_2_, 1mM Dithiothreitol and 5% glycerol) was added, making the concentration of DNA template to ~11.5ng/*μ*L. This mixture was then aliquoted into two tubes with 109*μ*L in each. Appropriate amount of MutS was added into each of the tubes so that the final MutS concentration was 950nM and 1900nM, respectively. The mixtures were then incubated at room temperature for 20 minutes. Equal volumes of Amylose Resin (New England Biolabs), washed and pre-equilibrated with 1X binding buffer, were added into the tubes. The mixtures were incubated at room temperature for 30 minutes, before being spun down. The supernatants were collected and MinEluted in 10*μ*L EB. We used 2*μ*L of the 1:100 diluted elution as the templates for the recovery PCR. Lastly, we pooled the PCR products, cleaned them up, and used them for the next iteration of MutS treatments.

### Next-Generation Sequencing using Illumina MiSeq

Each of the control and enzymatically treated samples was prepared as an individual sequencing library. In summary, the sequencing libraries were prepared using two rounds of real-time PCR, with the first round appending the Illumina P5 sequence and the second appending the P7 sequence as well as the indices. Specifically, the first round of PCR was set up by mixing 25*μ*L KAPA SYBR FAST Universal 2X qPCR Master Mix (KAPA Biosystems), 1*μ*L 10*μ*M Multiplexing PCR Primer 1.0, 1*μ*L 10*μ*M Multiplexing PCR Primer 2.0, 1*μ*L ∼100pg/*μ*L error correction DNA template, and 22*μ*L nuclease-free water. Per the manufacturer’s instructions, the 2-step thermocycling protocol was used for the real-time PCR reactions. Once the signals reached the plateaus, the reactions were stopped and cleaned-up using Agencourt AMPure beads, according to the manufacturer’s instructions. The final elution volume was 30*μ*L. To set up the second round of PCR, 25*μ*L KAPA SYBR FAST Universal 2X qPCR Master Mix, 1*μ*L 10*μ*M Multiplexing PCR Primer 1.0, 1*μ*L 10*μ*M PCR Primer each with a distinct index, 1*μ*L ∼100pg/*μ*L template from the first round PCR, and 22*μ*L nuclease-free water. The thermocycling and cleaned-up procedures remained the same as those in the first round of PCR. Then, the individually prepared sequencing libraries were quantified using the Library Quantification Kit-Illumina (KAPA Biosystems), according to the provided protocol. Barcoded libraries were subsequently mixed to ~10nM concentration, and the mixed libraries were quantified again before being loaded into MiSeq.

## Results

### Next-generation Sequencing Based Analysis of a Model Gene Assembly

To assess different enzymatic error correction methods, we first constructed a constant reference sequence that served as the base for downstream analyses. We designed this sequence to have (1) a length of 100 bp (not including two 21 bp priming regions for amplification and sequencing), (2) balanced nucleotide content (26:23:23:28 A:C:G:T content), (3) good coverage of all nucleotide pairs and most triplets (80%) while limiting homo-polymer repeats greater than two, and (4) a 28 bp region in the center that has good melting temperature and low secondary structure to facilitate overlap-extension assembly of the two primers. We assembled this sequence from two 85 nt oligos by a preliminary round of polymerase chain assembly (PCA). We then diluted the products of that reaction and used PCR to amplify the full length 142 bp construct (Figure 1). After assembly, we subjected the full-length products to two rounds of ten enzymatic error correction treatments and sequenced aliquots from the resulting reactions.

**Figure 1:**
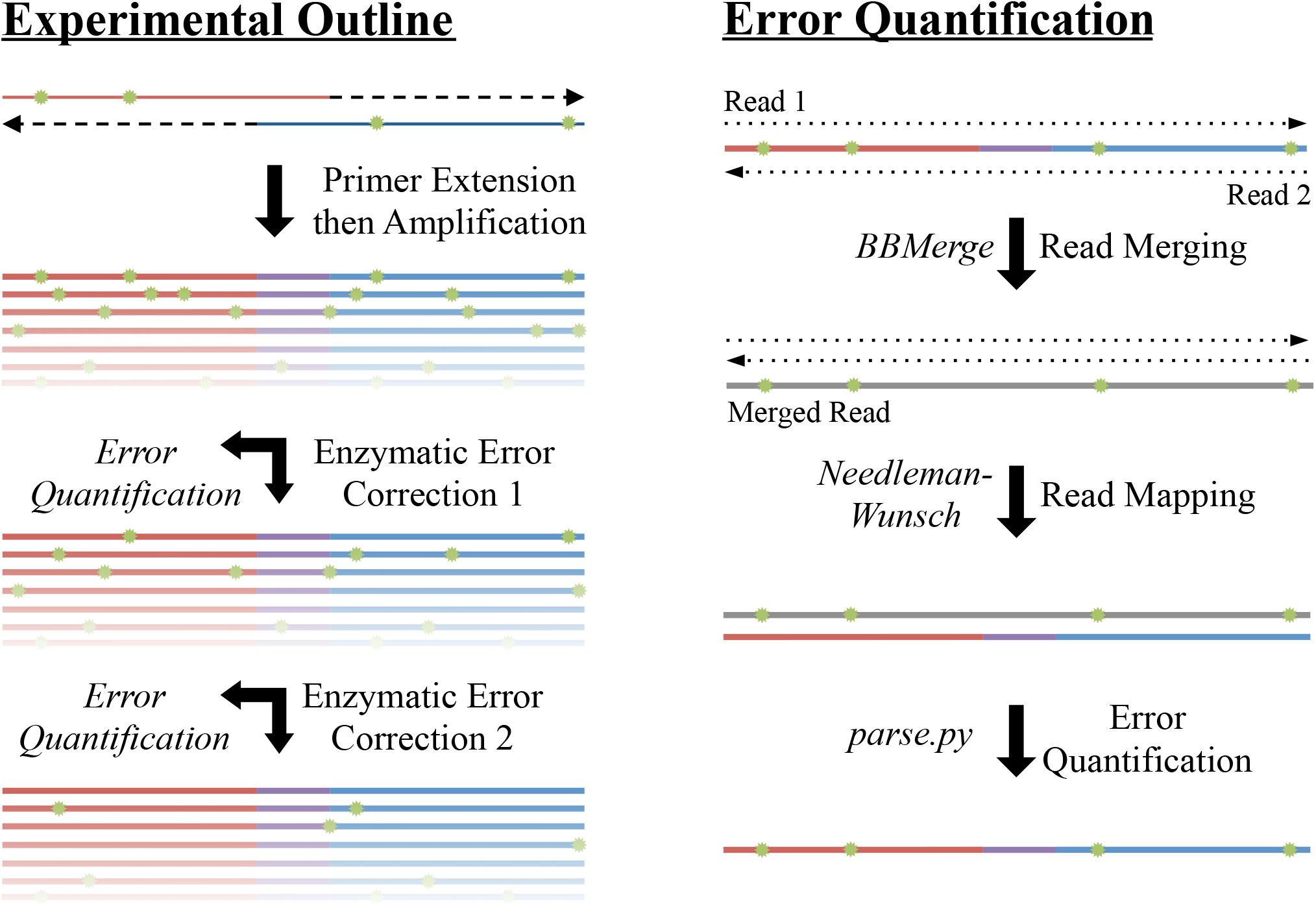
Schematic of Enzymatic Error Correction and Downstream Data Processing. We generated a template to subsequently error-correct, by annealing and primer extension, and then amplified via PCR. We then denatured and re-annealed the PCR products to form heteroduplexes, thereby exposing any errors (shown in green). We subjected the pool of heteroduplexes to two successive rounds of ten different enzymatic error correction treatments. We took aliquots from each round and sequenced the products on an Illumina MiSeq with fully overlapping forward and reverse reads. To mitigate sequencing errors, we used `BBMerge` to merge reads with a perfect agreement between the forward and reverse reads. We then aligned these sequences to the designed reference using an exhaustive Neeleman-Wunsch aligner to minimize alignment artifacts. Finally, we further processed the alignments to quantitate the types and extent of different errors across all conditions.

We expect that errors arising during sequencing will convolute our true signal. In order to limit these errors as much as possible, we developed a stringent data processing pipeline briefly outlined as follows: First, we cleaned our raw sequencing reads (509,717 per sample on average) by trimming sequencing adapters, removing any reads containg “N” base calls (212 reads on average), and filtering out any reads that aligned to either the PhiX or *E. coli* genomes with `BBDuk` (822 reads on average). This ensures that any spurious reads will not contaminate our alignments and lead to false-positive error calls. Next, we merged our paired end reads together with `BBMerge`, only keeping alignments with perfect correspondence between the forward and reverse reads. Since we sequenced our assembly with fully overlapping reads, each base is effectively sequenced twice. We found that an average of 95.2% of all bases in the merged reads had a Phred33 score (Q) of 41 (∼ 1/12,600 chance of being miscalled), and 99.8% of all bases on average were above Q30 (1/1000 chance of being miscalled). It should also be noted that most bases were probably above Q41 as this is the default maximum Phred score for most read mergers to maintain backwards compatibility with legacy software. The merging step removed an average of 15.8% of input reads, resulting in an average of 426,514 reads per sample at the end of processing.

After pre-processing the reads, we used a Python implementation of the Needleman-Wunsch aligner, `uta-align`, to align our reads to the perfect reference sequence. We elected to use a Needleman-Wunsch aligner as it is guaranteed to converge on the optimal alignment for a given scoring system [44, 45]. In contrast, typical short read aligners such as `BWA` and `Bowtie2` do not offer such guarantees as they use heuristics to trade accuracy for speed [46, 47]. We find that these heuristics often result in sub-optimal alignments and miscategorization of error sub-types (Figure S1, Table S1).

### Analysis of a Two Oligo Assembly

We first applied our pipeline to quantify the different types of errors found in a two oligo assembly of standard (not error doped) oligos (Figure 2). We find that on average about one-third of assemblies contain errors, with an overall error frequency of approximately 0.0047 errors per bp. We find that mismatches account for the majority of errors (∼75%), followed by single (∼14%) and multiple-base deletions deletions (~8%) (Figure 2A). The mismatches segregate into two significantly different populations, with the median error rate per base being higher at A’s (4.33 ×10^−3^) and T’s (4.25 × 10^−3^) than at G’s (1.68 ∼ 10^−3^) and C’s (1.91 ∼ 10^−3^) (Figures 2B, C; Mann-Whitney U, *p* ≪ 0.001, Holm-corrected). Furthermore, we find that the median rate of transitions was significantly higher than that of transversions for each base (Figures 2C; Mann-Whitney U, *p* ≪ 0.001, Holm-corrected). All of these observations indicate that the mismatch error rate is due to polymerase misincorporation during the amplification steps for assembly and sample-preparation for sequencing. Specifically, we used Taq-based polymerase during next-generation sequencing library preparation steps, which is known to misincorporate bases at a higher frequency at A’s and T’s [48]. Consistent with our observations, the misincorporations caused by Taq are preferentially A/T → G/C transitions [49, 50]. Lastly, our error frequency of 0.0047 errors/bp is consistent with approximately 30 rounds of amplification with a polymerase error rate of ~ 2.5 ∼ 10^−4^, which is within the range of previously reported Taq error rates [51, 52].

**Figure 2:**
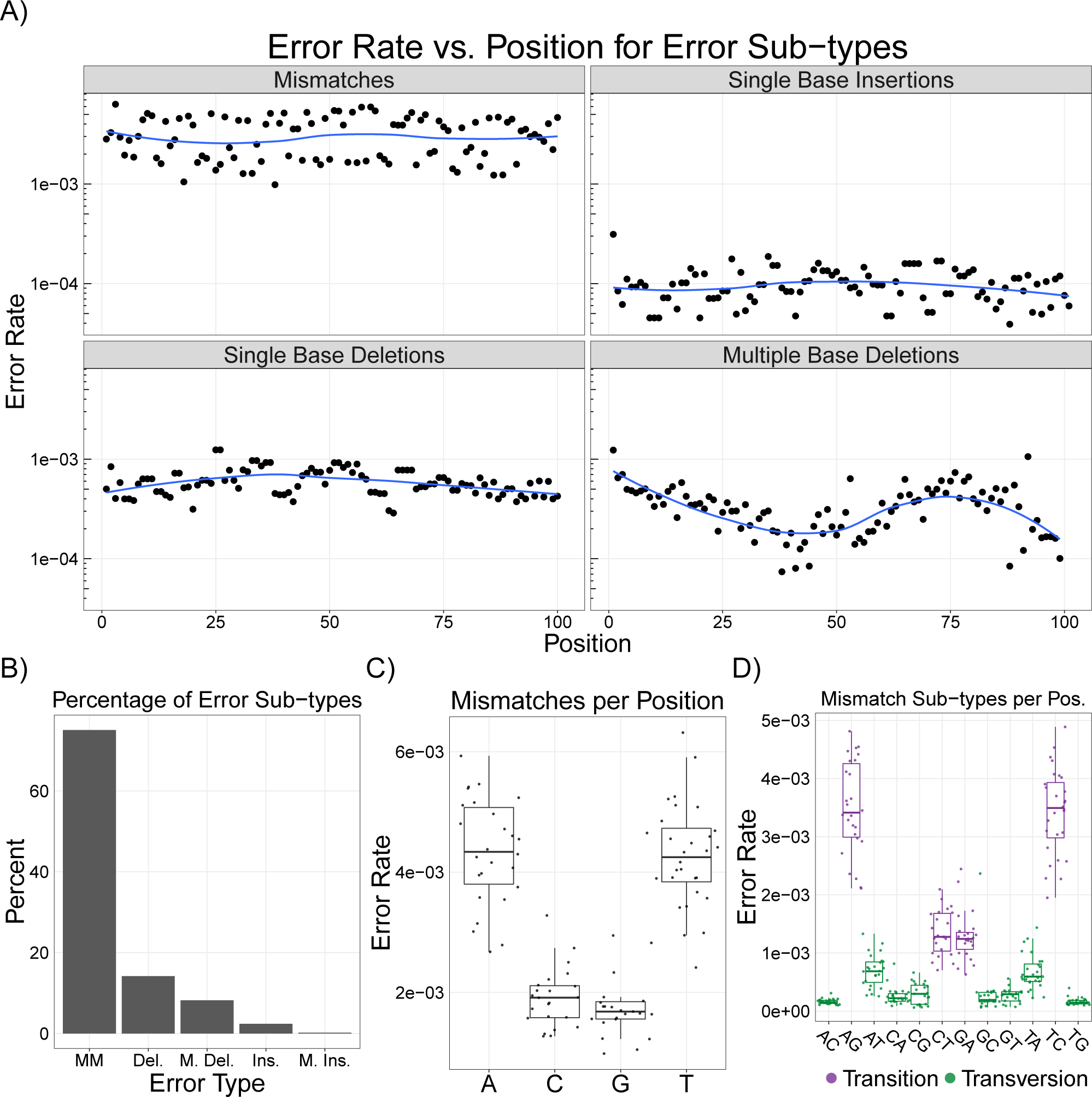
Analysis of Model Gene Assembly Error Rates. (**A**) The error rates per base are plotted across each position in our model separated by the four major classes of error types. We do not see strong positional effects for errors across the template. (**B**) We find a majority of errors on the template are mismatches (MM), followed by single (Del.) and multiple base (M. Del.) deletions; Single (Ins.) and multiple base (M. Ins.) insertions occur at even lower frequencies. (**C**) We measure a significantly higher mismatch rate at A’s (4.33 × 10^−3^) and T’s (4.25 × 10^−3^) than at G’s (1.68 × 10^−3^) and C’s (1.91 × 10^−3^) (Mann-Whitney U, *p* ≪ 0.001). (**D**) We measure a significantly higher number of transitions (red) than transversions (blue) at each base (Mann-Whitney U, *p* ≪ 0.001). The higher error rates at A’s and T’s is consistent with Taq polymerase errors. **Note:** Blue line is a LOESS fit; box plots are first and third quartile for hinges, median for bar, and 1.5 × the inter-quartile range for whiskers.

Next, we quantified the rates of single- and multiple-base deletions. We find that the median single-base deletion rate per position (5.64 × 10^−4^), and that this rate did not vary significantly over the positions (Mann-Whitney U, NS). We also find that multiple-base deletions occur at a similar rate as single-base deletions (3.35 × 10^−4^), and measure positional effects for where they occur. Some of this dependence can be explained by the fact that the position of multiple-base deletions are mapped to the left-most deleted base. Thus, we expect the total number of multiple-base deletions to be highest at position one and decreasing after, since there are the most possible combinations of multiple-base deletions at that position. In addition, we measure a significant decrease in the median multiple-base deletion rate in the annealing region (positions 36-64) of our assembly (Mann-Whitney U, *p* ≪ 0.001). Large deletions in this region would disrupt the hybridization of the initial assembly, leading to sequence drop-outs and a decrease in the measured number of deletions. We also expect the multiple deletion rate to drop towards the end of the sequence due to a “TATATAT” motif at positions 92-98. Any “TA” deletion (or other substring contained multiple times in the motif) will map to the left-most position, 92.

Finally, we quantified single-base insertions. These errors occur at median rate per position of 9.65 × 10^−5^ and exhibit no positional dependence besides an outlier at position 1. This outlier can be explained by an incomplete primer trimming by `BBDuk`. Here, 57 of the 152 single-base insertions are a “T,” corresponding to the last base of the primer sequence directly upstream of our first base. Without these 57 bases, the rate of single-base insertions falls closer to the expected median value. Our method is also able to detect multiple-base insertions, which occur at a median rate of 6.16 × 10^−6^, but we omit them from the Figure 2A due to their rarity.

### Error-doped Oligos Enable Comparisons

In order to assess the sensitivity of our assay, we treated the standard oligo product with the error correction cocktail ErrASE and measured the resulting error rates (Figure S2). Although we were able to measure significant (Mann-Whitney U, *p* ≪ 0.001, Holm-corrected) reductions in the rate of single-base deletions, multiple-base deletions, and single-base insertions, we were not able to find a significant (Mann-Whitney U, NS, Holm-corrected) reduction between the median error rate of mismatches. As we discuss above, the significant amount of PCR required to prepare samples for sequencing introduces mismatch errors that mask the effect of error correction.

To ensure that we had a measurable change in error rates for mismatches after enzymatic treatment, we assembled our template from oligos that had errors doped into the sequence. Specifically, we ordered each base with 97% of the intended base, and 1% of the other three nucleotides (not including the 21 bp priming region and the last base of the oligo). Our measured mismatch frequency of ∼0.04 errors per bp was slightly higher than the designed 0.03 errors per bp. This error-enriched template allows us to accurately assess error-correction by enzymatic processes by increasing the rate of mismatches at an order of magnitude above limit of detection established for the standard oligo assembly.

As in Figure 2, we analyzed error rates of the resulting assembly from these error-doped oligos (Figure S3). Unlike the standard oligo assembly, we found no significant difference between the median mismatch rate (3.99 × 10^−2^) at any of the four bases (Mann-Whitney U, NS). Similarly, the median rate of individual transitions and transversions were not significantly different from each other (Mann-Whitney U, NS). These data suggest that incorrect bases were doped in to our oligos at an approximately equal rate that exceeded the baseline error rate of our polymerases. Unlike the standard assembly, we were unable to detect any positional biases for multi-base deletions. Lastly, we found the median error rate of all error sub-types to be higher in the error-doped assembly (Table 1, Figure S4; Mann-Whitney U, *p* ≪ 0.001). Although this is expected for mismatches, we suspect that the higher median error rates for the other error sub-types are a result of the non-standard synthesis required to dope the errors into our oligos.

**Table 1:** **Median error rate per position for assemblies using the error-doped oligos or the standard oligos.** We measure significant (Mann-Whitney U, *p* ≪ 0.001) differences between the median error rates of the error-doped and standard oligos for all error sub-types.

| Error Type | Doped Oligos | Standard Oligos |
| --- | --- | --- |
| All Errors | $4.38 \times 10^{-2}$ | $4.18 \times 10^{-3}$ |
| Mismatches | $3.99 \times 10^{-2}$ | $3.08 \times 10^{-3}$ |
| Single Deletions | $1.28 \times 10^{-3}$ | $5.64 \times 10^{-4}$ |
| Multiple Deletions | $1.17 \times 10^{-3}$ | $3.35 \times 10^{-4}$ |
| Single Insertions | $7.64 \times 10^{-4}$ | $9.65 \times 10^{-5}$ |
| Multiple Insertions | $6.16 \times 10^{-4}$ | $6.16 \times 10^{-6}$ |

### Enzymatic Error Correction Improves Assembly Quality

Having established the error profile of the error-doped assembly, we evaluated 10 different enzymatic error correction methods using six different enzymes on their ability improve the quality of this assembly (Figure 3). As expected, consecutive rounds of enzymatic error correction improved both the relative error frequencies and the number of perfect assemblies. ErrASE was the most effective at decreasing the error frequency, with two rounds of treatment dropping the error frequency from the doped oligo rate of 46.1 to 8.0 errors/kb. The next most effective enzyme at decreasing error frequency was T7 Endonuclease I (9.1 errors/kb). Based on previous reports in the mutation detection literature, we hypothesized that the addition of a ligase with T7 Endonucluase I would improve correction [39]. We find that the addition of T7 ligase actually decreased assembly quality relative to the no ligase control. In agreement with previous studies, we also find T7 Endonuclase I to be highly sensitive to protocol and concentration as exhibited by the wide range of error frequencies [12, 14]. After T7 Endonuclease I, we found MutS to be the third most effective enzyme at 11.2 errors/kb, with T4 Endonuclease VII, Surveyor, and Endonuclease V following.

**Figure 3:**
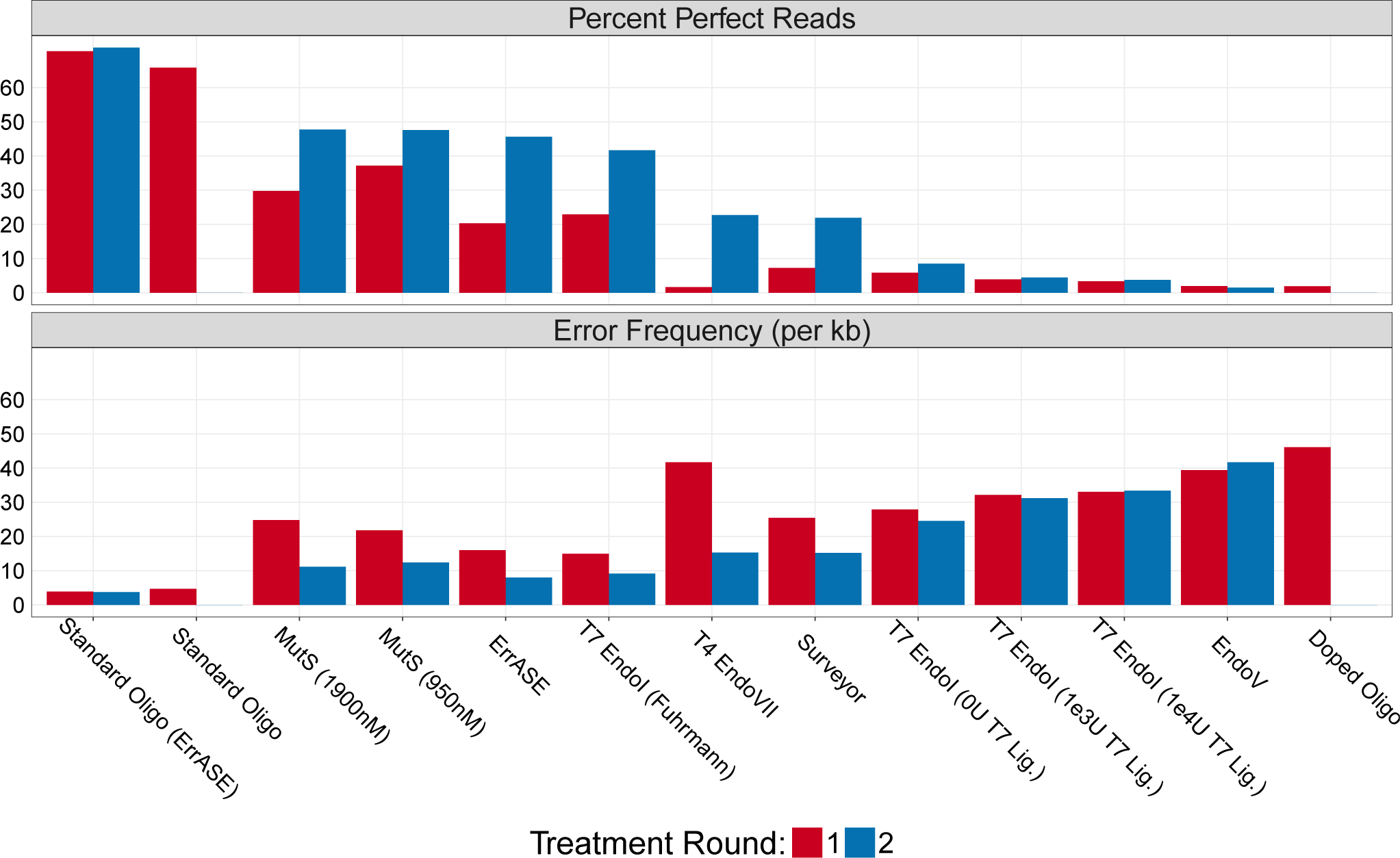
Effectiveness of Enzymatic Error Correction Methods. Here we compare the error frequency (errors/kb) and number of perfect assemblies for ten different enzymatic error correction methods. We find that MutS is the most effective enzyme at increasing the percentage of perfect assemblies. However, ErrASE is the most effective at decreasing error frequency. Additionally, we see that the efficacy of T7 Endonuclease I is dependent on protocol, and that the addition of a ligase had detrimental effects on sequence quality. **Note:** the x-axis is ordered by decreasing number of perfect assemblies.

However, when looking at number of perfect assemblies sequences, MutS was the most effective enzyme treatment. MutS increased the percentage of perfect sequences in the doped oligo from 1.9% to 47.8% (47.6% for 950nM), while ErrASE increased it to 45.6%, and T7 Endonuclease I increased it to 41.7%. In other words, the oligos that are imperfect after the MutS treatment have more errors on average than those after the T7 Endonuclease I and ErrASE treatments.

### Differences in Enzymatic Error Correction

With an average of 426,514 reads per round of error correction, our method provides sample sizes three to four orders of magnitude higher than any previous study. This enabled us to compare the effectiveness of these enzymes on rarer errors such as insertions that would be inadequately sampled with Sanger sequencing. Using the error-doped template as a reference, we measured the relative change in error rates for each position across all different enzymatic error correction methods (Figure 4).

**Figure 4:**
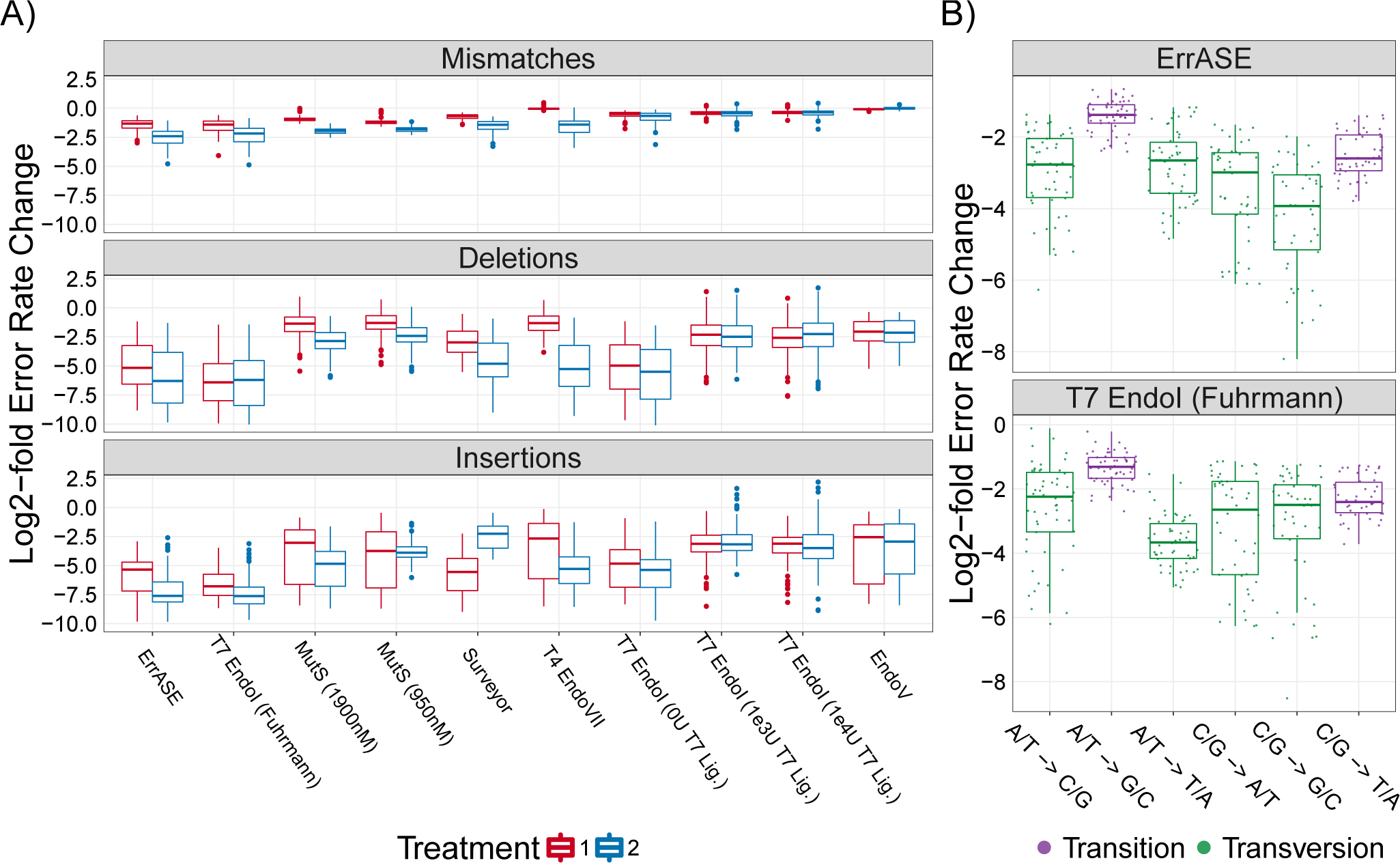
Relative Decrease of Different Error Types. (**A**) All enzymes were able to correct both single- and multiple-base insertions and deletions. Additionally, we find that the best performing enzymes corrected the highest amount of mismatches. **Note:** the x-axis is ordered by increasing error frequency. (**B**) We measure significant differences between the median decrease in C/G → G/C mismatches and the bulk median of all other mismatches after two treatments of ErrASE. Similarly, two treatments of T7 Endonuclease I results in a significant difference between the median decrease in A/T → T/A mismatches compared to the bulk median of all other mismatches (both Mann-Whitney U, *p* ≪ 0.001).

We see that in general, all enzymes tested were able to correct insertions and deletions. We find that enzyme performance (as measured by error frequency or number of perfect assembliess) is directly related to the ability to correct mismatches. For example the best performing enzymes, ErrASE, T7 Endonuclease I, and MutS, were able to decrease the median mismatch error rate relative to the error-doped input by 6.2-, 5.1-, and 4.2-fold, respectively. In contrast, the worst performing enzyme, Endonuclease V, was unable to decrease the median mismatch error rate relative to the error-doped input. We note that these measurements represent a ceiling on mismatch correction, as it is likely that the polymerase added mismatches back into the sample during the sequencing preparation.

We next sought to measure differences in affinity for specific errors between enzymes (Figures S5–S7). We were unable to measure any significant differences between bases for the median fold reduction of insertions and deletions (Kruskal-Wallis, NS) across all enzymes after two treatments. However, we were able to detect significant differences between the median fold reduction of different mismatches (Kruskal-Wallis, *p* ≪ 0.001) across all enzymes after two treatments. Based on these data, we searched for specific mismatch correction biases in our best performing enzymes. For example, we found that two rounds of ErrASE or MutS treatment resulted in a significantly different change in the median fold reduction of C/G → G/C mismatches as compared to the bulk median of all other mismatches (15.2- vs 5.4-fold for ErrASE; 5.1- vs 4.1-fold for MutS; Mann-Whitney U, *p* ≪ 0.001). In contrast, two rounds T7 Endonuclease I did not result in significant changes in the median fold reduction of C/G → G/C mismatches (5.6- vs 5.1-fold; Mann-Whitney U, NS). They did however, significantly change the median fold reduction of A/T → T/A mismatches as compared to the bulk median of all other mismatches (12.7- vs 4.2-fold; Mann-Whitney U, *p* ≪ 0.001). Lastly, we found that using a T7 Ligase in conjunction with T7 Endonuclase I actually decreased assembly quality relative to the no ligase control, in contrast to previous reports [39]. These data also suggest that T7 Endonuclase I is sensitive to protocol, as the no ligase control performed worse than the Fuhrmann *et al.* protocol [12].

Taken together, these data suggest that different enzymatic error correction methods could be used for different applications. For example, GC or AT rich constructs would be best corrected by ErrASE and T7 Endonuclease I, respectively. Alternatively, MutS can be used for applications such as protein libraries, where the proportion of perfect sequences are paramount. We also note that the relative rate of correction for transitions is likely lower than what is measured here due to polymerase errors. For example, the median fold correction of A/T → G/C transitions (the most common Taq-based error) was significantly different than that of the bulk median for all other mismatches for ErrASE, MutS, and T7 Endonuclase I (2.6- vs 7.1-fold for ErrASE; 2.8- vs 4.4-fold for MutS; 2.5- vs 6.8-fold for T7 Endonuclease I; Mann-Whitney U, *p* ≪ 0.001).

## Discussion

One of the most promising methods to improve the quality of gene synthesis products is enzymatic error correction. Previous characterizations of error correction enzymes were limited by Sanger sequencing, which prohibited deep enough sequencing to adequately sample rare variants. Here we surpass this bottleneck by leveraging next-generation sequencing (NGS) and a custom computational pipeline to analyze errors in a model gene assembly. With sample sizes of three to four orders of magnitude greater than any previous study, we were able to accurately sample rarer errors such as insertions. In addition, NGS precludes the need for time consuming cloning steps. This enabled us to rapidly compare six of the most commonly used error correction enzymes in a total of eleven different conditions in a single experiment, and marks the first comprehensive comparison of enzymatic error correction methods via NGS.

We took multiple steps to minimize the number of false error calls resulting from our method. First, we sequenced our assembly with fully overlapping paired-end reads. Since each base is called independently twice and we only merge reads with a perfect match between the forward and reverse reads, it is unlikely that a large number of sequencing errors made it through this filter. Next, we compared the error profile of the Needleman-Wunsch alignment to two commonly used short-read aligners, `BBMap` and `Bowtie2`. As `BBMap` and `Bowtie2` use heuristics that trade accuracy for speed, we found that their resulting alignments were sub-optimal and led to higher false error calls relative to the Needleman-Wunsch alignment.

With our method tuned, we then analyzed errors in model a two-oligo assembly to assess its sensitivity. Previous studies found the most common errors in the final gene synthesis product to be either single-base deletions [11, 14, 19], or mismatches [12, 20]. Here, we also found mismatches to be the most common error in our assembly. However, we believe that the majority of these mismatches were introduced by our Taq-based polymerase during the NGS library preparation steps by comparing the specific mismatch rates to those previously reported in the literature, [48, 49, 50, 51, 52]. In order to ensure that we could measure changes in the amount of mismatches, we re-assembled our model sequence with oligos synthesized with 3% of the incorrect base at every position. We expected that the net change in mismatches in the error-doped template after error correction would be larger than the basal error rate of the polymerase, enabling quantification. Additionally, increasing the error rate gives a more realistic number of errors (3-4) per assembly that might occur in a longer gene synthesis.

We then used our method to test the ability of six of the most common error correction enzymes in eleven total conditions to improve the quality of the error-doped assembly. As expected, we found that the all of the error correction enzymes were able to decrease the error frequency and increase the number of perfect assemblies. We also found that two consecutive treatments of error correction were more effective than one, although further rounds of treatment will likely yield diminishing returns with respect to the level of error correction. We then leveraged the large sample sizes generated by NGS to probe specific differences between different enzyme treatments. These data suggest that ErrASE would bet the most effective at correcting GC-rich templates, and T7 Endonuclease I is the most effective at correcting AT-rich templates. Alternatively, MutS would be appropriate for the most common applications requiring a single sequenced-verified perfect assembly. The discrepancy of average error frequency and percentage of perfect sequences highlights of using the metrics that are most appropriate for downstream application. In addition, we find that performance of these enzymatic treatments is sensitive to the protocol used as shown in the MutS and T7 Endonuclease I assays.

Our method in its current iteration is not without limitations. For one, any polymerase misincorporations will convolute the true mismatch correction rate of a a given enzyme. This issue can be partially ameliorated by using a high-fidelity polymerase throughout the assembly and NGS library preparation steps. Alternatively, we can incorporate random barcoding strategies or utilize single molecule sequencing to further eliminate polymerase errors [49, 52]. Second, Illumina sequencing limits our assessments to assemblies < 600 bp. We could extend our methodology to long-read technologies such as PacBio or Oxford Nanopore to assess kilobase-scale gene synthesis products [53]. At these lengths, we would likely have to switch from a Needleman-Wunsch alignment to more optimized versions in order to avoid a significant time penalty [54]. Lastly, our model assembly is not indicative of a typical gene synthesis product as it does not code for a gene, is shorter than standard assemblies (142), is assembled from only two oligos, and has a contrived mismatch error rate. The artificial source of errors is especially important, as depending upon protocol, oligo source, assembly conditions, and use case, both the input and desired error profiles may change.

Overall, our method is a fast and accurate method for looking at errors in arbitrary sequences. We believe that this method will be useful for not only rapidly profiling new enzymatic error correction methods, but for other applications such as assessing the quality of chip-synthesized oligos or developing new gene synthesis methods.

## Availability

The computational pipeline described above is open source, free to use under the MIT license, and available at https://github.com/kosurilab/errorCorrect. For the final analysis and figure production, we used R (version 3.3.*) and ggplot2 [55, 56].

## Accession Numbers

Sequencing data are available from the sequencing read archive (SRA) with the accession number XXXXXXXX [pending].

## Acknowledgments

The authors would like to acknowledge members of the Kosuri Lab for comments on the manuscript, especially Rocky Cheung and Calin Plesa.

---

**Figure S1:**
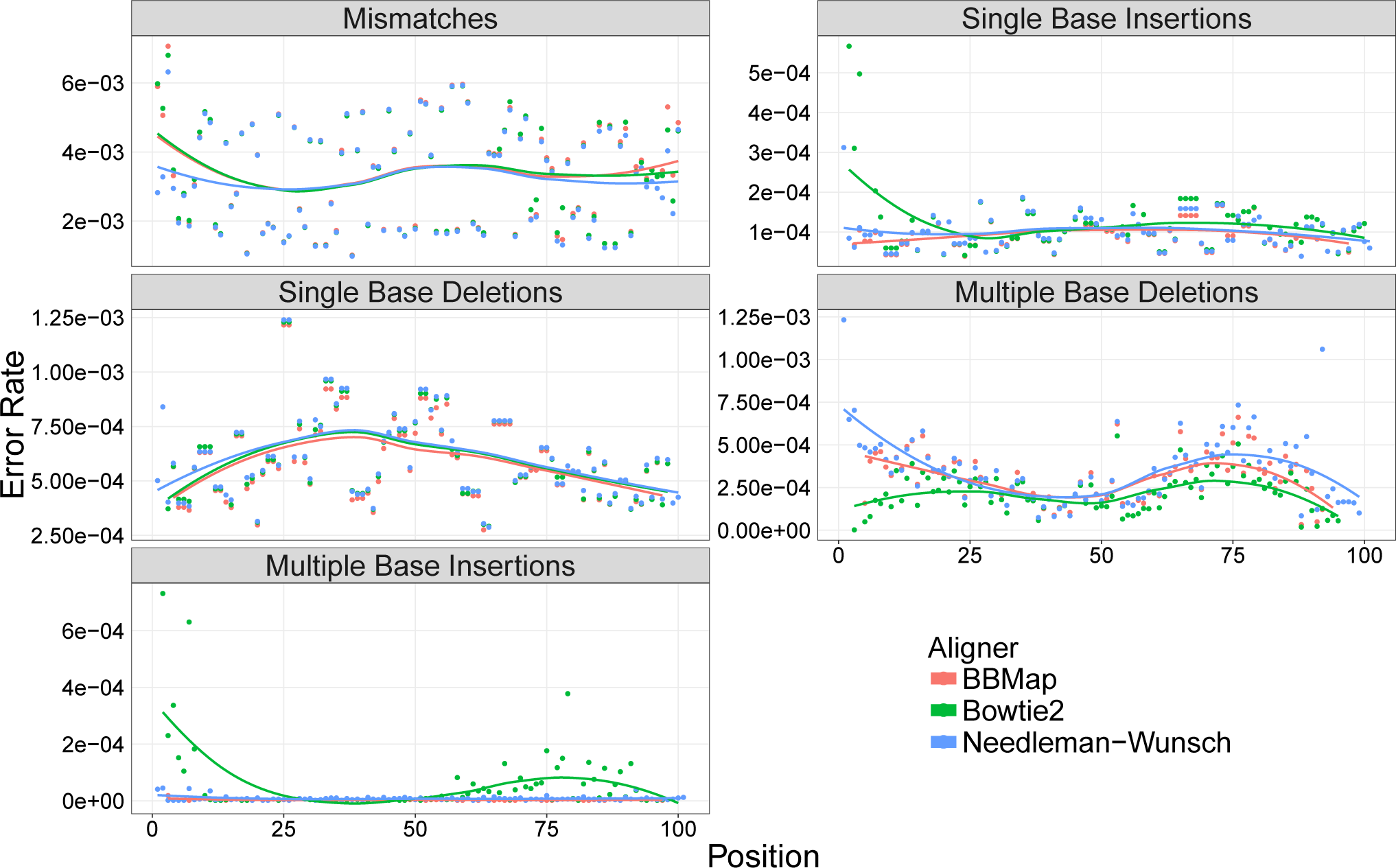
Effect of read aligner on error rates. Here we mapped reads from the standard IDT oligo with `BBMap` (red), `Bowtie2` (green), and our Needleman-Wunsch aligner (blue), and quantified the error rates with our pipeline. We see that the choice of aligner affects the resulting error rates, especially for detecting multiple-base deletions.

**Figure S2:**
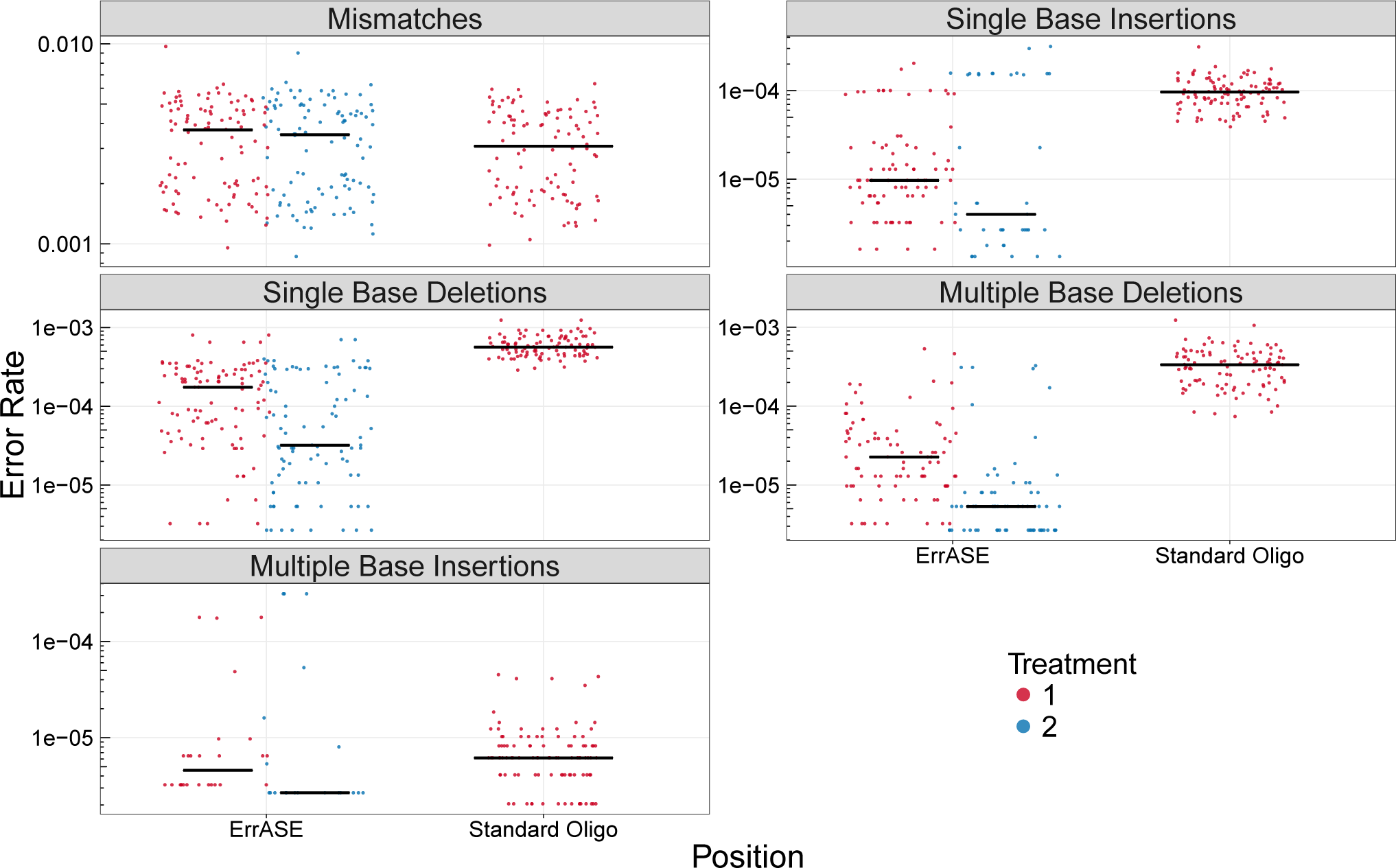
Distributions of error rates per position for the standard oligo assembly before and after ErrASE treatment. We were unable to detect a significant change between the median error rate after two treatments for mismatches. **Note:** black bar is median value.

**Figure S3:**
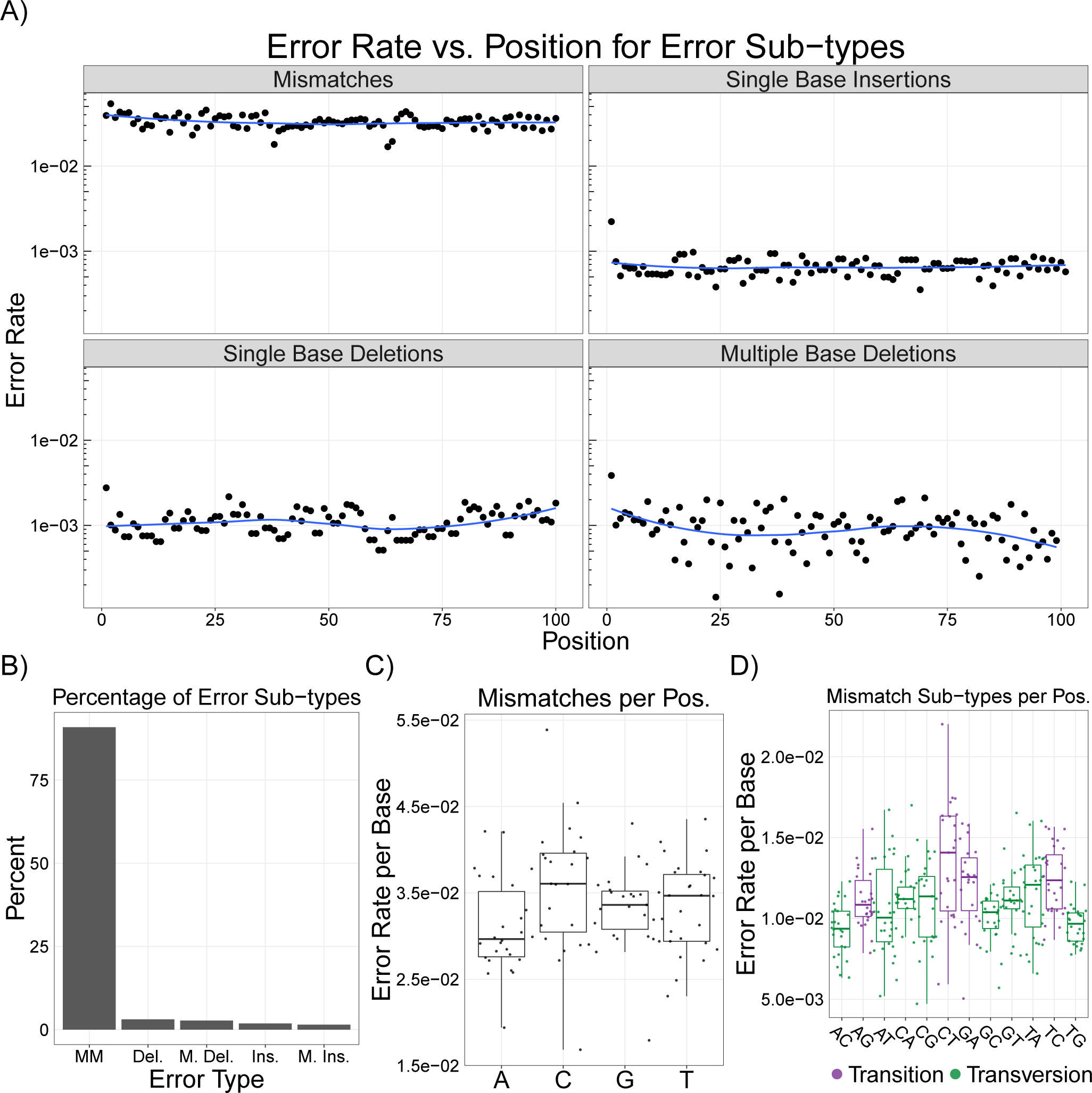
In-depth analysis of error-doped assemblies. Here, we find no significant difference in the median mismatch rate of all four bases, median transition or transversion rate, or rate of single-base deletions for each base (all tests were Mann-Whitney U, NS, Holm-corrected). **Note:** here we performed the same analysis as Figure 2 in the main text with the error-doped assembly.

**Figure S4:**
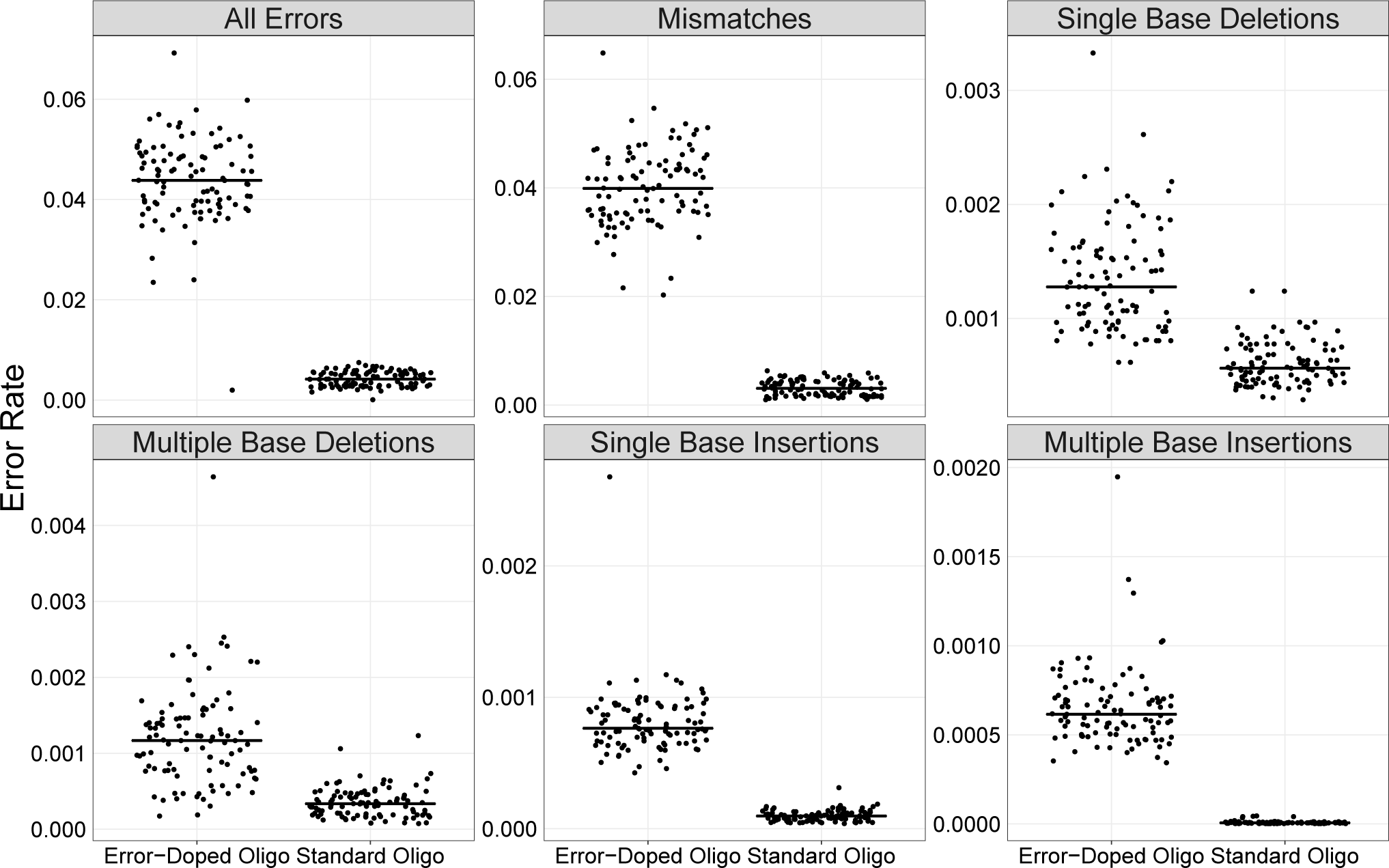
Comparison of measured error rates from error-doped and standard oligos. Here we plot the distribution of error rates per position and see that for every error sub-type the error rates are significantly higher for the error-doped oligos than those produced by the standard process (Mann-Whitney U Test, all *p* ≪ 0.001). **Note:** Black bar is the median value.

**Figure S5:**
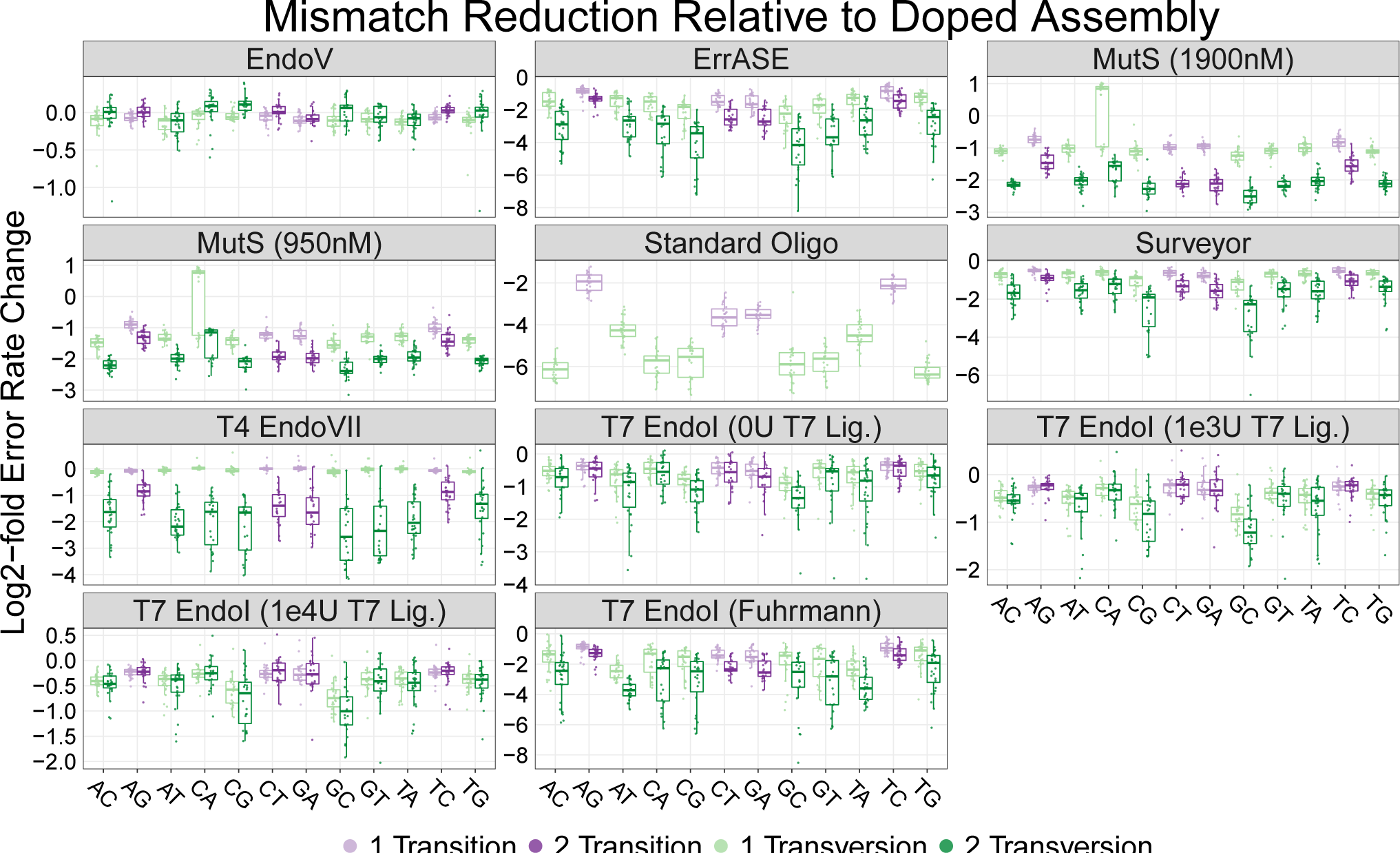
Mismatch correction preferences relative to the error-doped oligo for every enzyme across two consecutive treatments. Error rates are plotted as the log_2_-fold-change in error rate relative to the error-doped template. **Note:** box plots are first and third quartile for hinges, median for bar, and 1.5× the inter-quartile range for whiskers.

**Figure S6:**
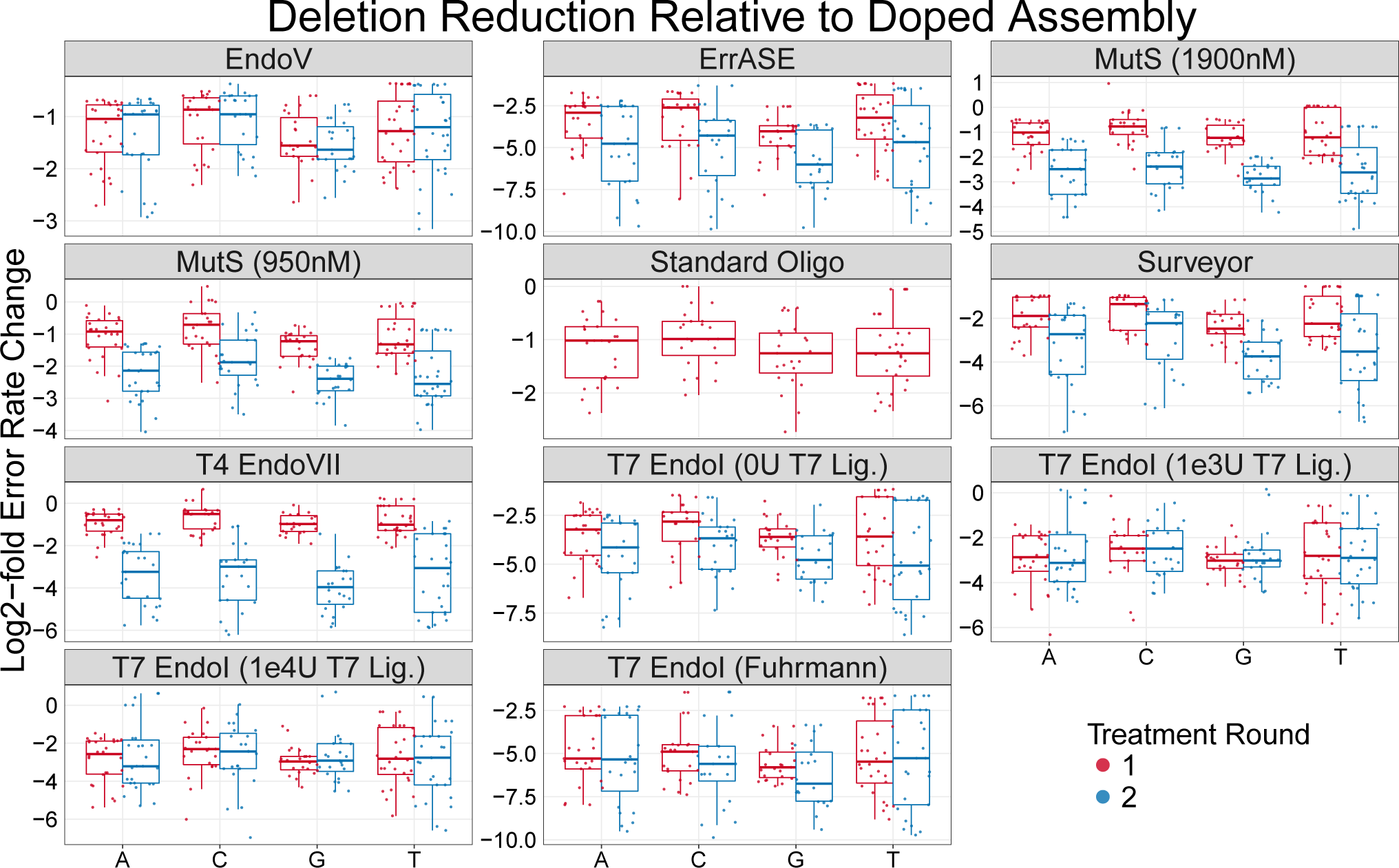
Single-base deletion correction preferences relative to the error-doped oligo for every enzyme across two consecutive treatments. Error rates are plotted as the log_2_-fold-change in error rate relative to the error-doped template. **Note:** box plots are first and third quartile for hinges, median for bar, and 1.5× the inter-quartile range for whiskers.

**Figure S7:**
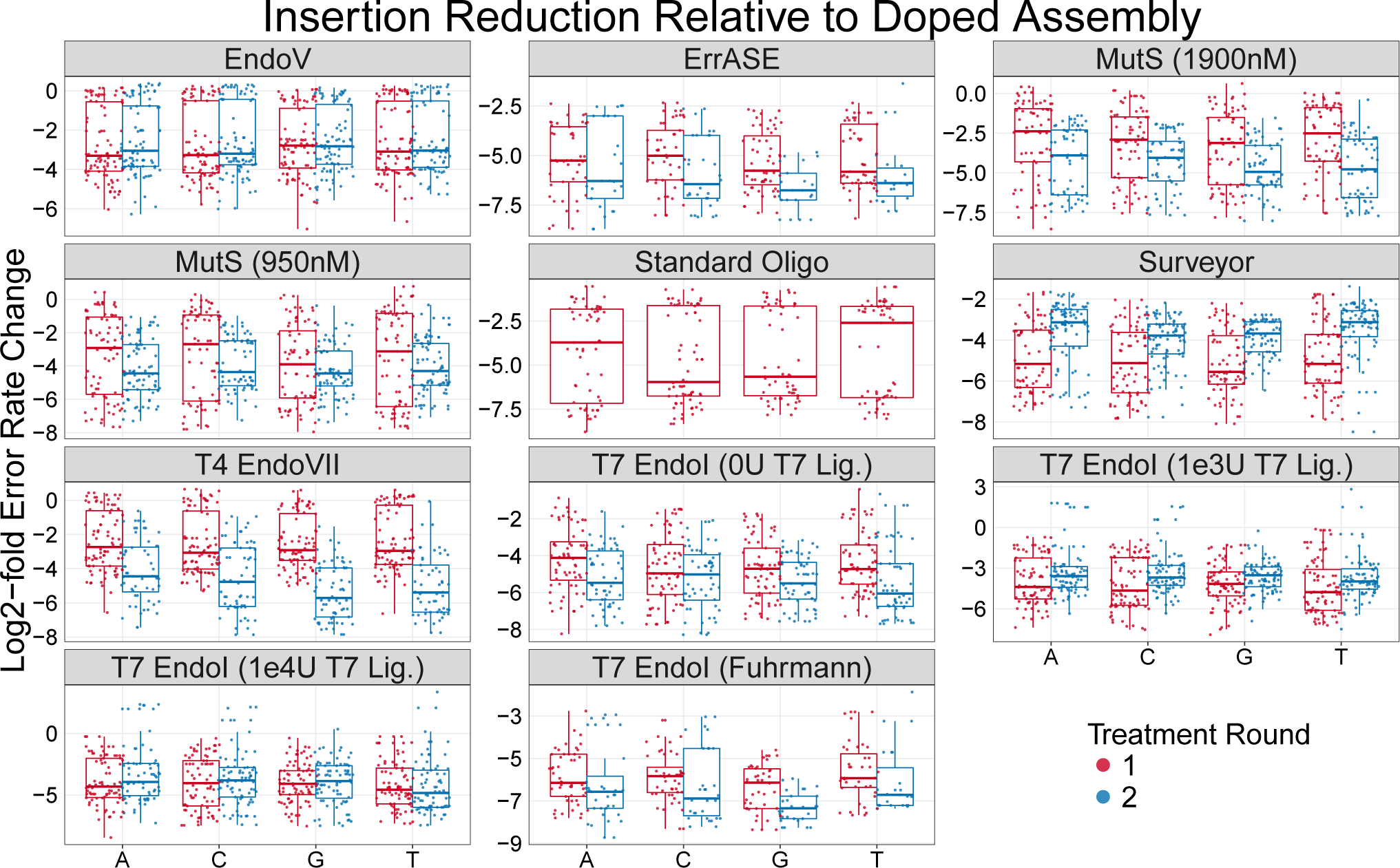
Single-base insertion correction preferences relative to the error-doped oligo for every enzyme across two consecutive treatments. Error rates are plotted as the log_2_-fold-change in error rate relative to the error-doped template. **Note:** box plots are first and third quartile for hinges, median for bar, and 1.5× the inter-quartile range for whiskers.

**Table SI:**
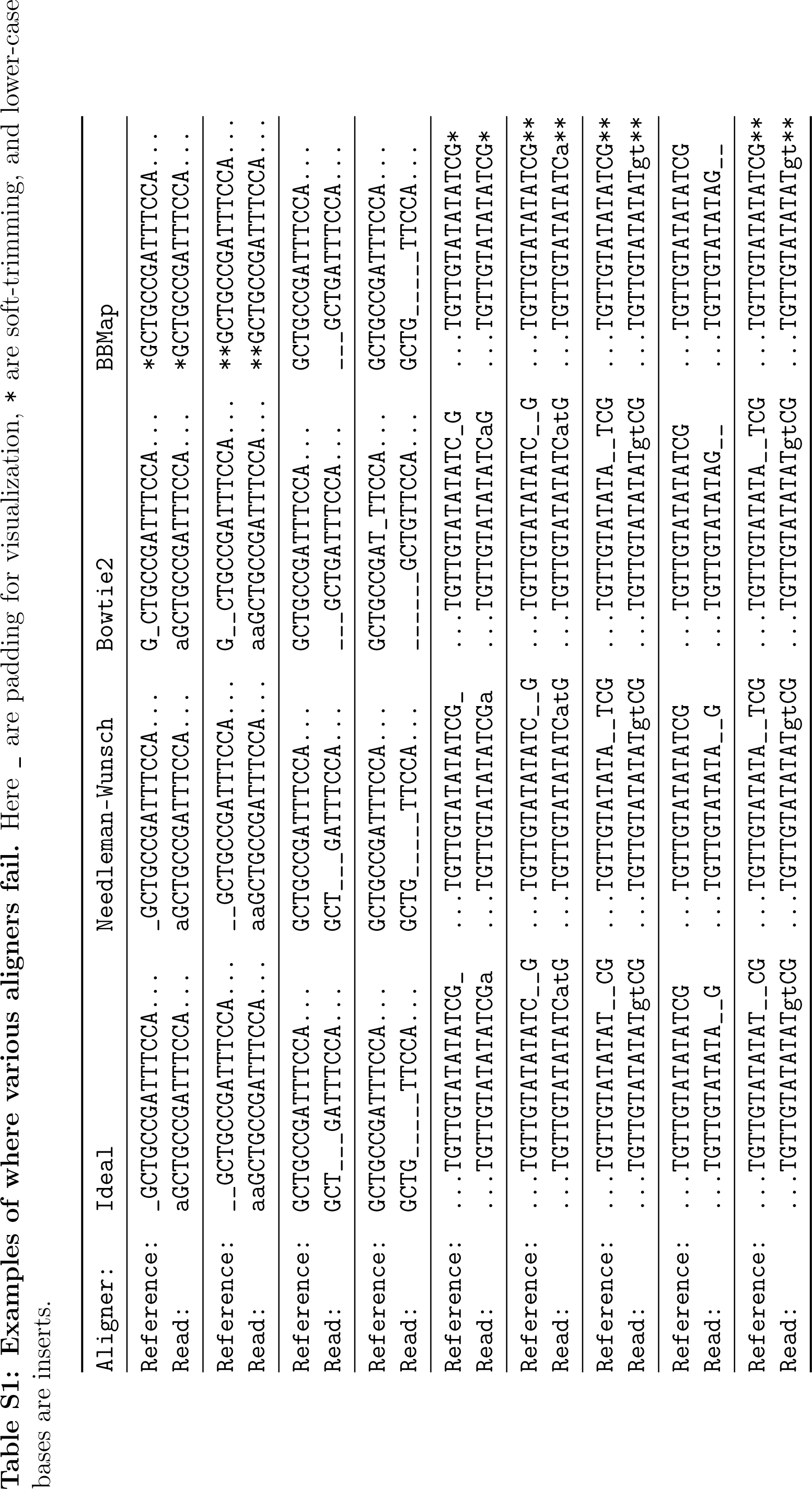
**Examples of where various aligners fail.** Here _ are padding for visualization, * are soft-trimming, and lower-case bases are inserts.

